# Plasma proteome screen of individuals with elevated blood counts to identify diagnostic signatures and biomarkers inherent to patients with myeloproliferative neoplasms

**DOI:** 10.64898/2026.08.03.742450

**Authors:** Xiangfu Zhong, Ingrid Lundahl, Axel Rosell, Roza Chaireti, Johanna Ungerstedt

## Abstract

**Background:** The Philadelphia negative myeloproliferative neoplasms (MPN), including essential thrombocythemia (ET), polycythemia vera (PV) and primary myelofibrosis (PMF), are characterized by myeloid cell proliferation, thrombosis and inflammation. Suspicion of MPN arises from increased blood count in one or more lineages; however, knowledge on the MPN plasma proteome including biomarkers measurable in blood, are lacking. Comparing the plasma proteome of MPN patients to subjects with elevated blood counts but without MPN diagnosis, may provide an increased understanding of the MPN disease biology as well as diagnostic biomarkers measurable in blood.

**Patients and methods:** We performed plasma proteome profiling in 87 patients referred to the Department of Hematology due to elevated blood counts. Of these, 55 were diagnosed with MPN and 32 did not fulfill MPN diagnostic criteria and thus constituted the non-MPN control group.

**Results:** The frequency of thrombosis was equal between the groups. We found 189 differentially expressed proteins between MPN and non-MPN, enriched for Hemostasis and Platelet activation proteins. Using Lasso multiple regression, we identified SORT1, GP1BA, PSPN, MMP1 and BAG6 separating MPN from non-MPN individuals, and TFRC and SEMA7A specific for MPN subtype PV. Interestingly, TFRC alone had a diagnostic accuracy for identifying PV of 89.3%, and when combined with serum erythropoietin it increased to 99.3%. Only two proteins, IL-6 and GH1, were increased in *JAK2* mutant MPN compared to *JAK2* wildtype MPN. The same trend was seen for *JAK2* mutant ET compared to *JAK2* wildtype ET, indicating that IL-6 is induced by JAK STAT activation. However, IL-6 levels did not differ between MPN and non-MPN patients.

**Discussion/conclusion:** In conclusion, hemostasis and platelet activation are inherent to MPN disease whereas little difference was found in proinflammatory cytokines between MPN and non-MPN groups. We demonstrate novel potential blood biomarkers for MPN and MPN subtypes, in particular TFRC for identifying PV patients.

## Introduction

Philadelphia-negative myeloproliferative neoplasms (MPN) are a group of clonal hematopoietic stem cell disorders (Mead and Mullally 2017; Vainchenker and Kralovics 2017). The classical MPNs are essential thrombocythemia (ET), polycythemia vera (PV) and primary myelofibrosis (PMF). In recent years, it has been increasingly demonstrated that the driver mutations of MPN arise early in life (Williams et al. 2022); however, the median patient age at MPN diagnosis is around 70 years of age (Sud et al. 2018; Batyrbekova et al. 2025).

The three predominant driver mutations in MPN are *JAK2*, *CALR*, and *MPL* (James et al. 2005; Levine et al. 2005; Kralovics et al. 2005; Pikman et al. 2006; Baxter et al. 2005; Nangalia et al. 2013). *JAK2* V617F mutation is the most frequent driver mutation, encountered in >95% of PV (Kralovics et al. 2005; Levine et al. 2005; Baxter et al. 2005), and 50-60% of ET and PMF cases (Tefferi and Vainchenker 2011). *JAK2* exon 12 mutations are seen in the remaining 2-5% cases of PV (Chuang et al. 2025; Scott et al. 2007) but not in ET or PMF, whereas *MPL* mutation occurs in 5%-10% of ET and PMF, and *CALR* mutation in 25%-35% of all ET and PMF cases (Klampfl et al. 2013; Nangalia et al. 2013). Mutation analysis and bone marrow examination is required for MPN diagnosis (Khoury et al. 2022; Arber et al. 2022).

Suspicion of MPN is usually raised by elevated blood counts; however to date there are no specific laboratory biomarkers of MPN or MPN subtypes, with the exception for serum erythropoietin which is frequently suppressed in PV and included as a minor diagnostic criterion for PV (Arber et al. 2016; Szuber et al. 2018; Arber et al. 2022). Diagnosis therefore relies on bone marrow morphology, and molecular testing for driver mutations, none of which is pathognomonic. This highlights the need for additional biomarkers, preferably blood-based for easier access and availability, that can improve diagnostic precision by avoiding unnecessary bone marrow sampling, distinguish MPN subtypes, and capture disease-associated biology beyond blood cell counts.

Patients with MPN have an elevated risk of arterial and venous thromboembolism, with the highest risk within the first 2 years after diagnosis and an overall lifetime risk around 20% (Batyrbekova et al. 2025; Rungjirajittranon et al. 2019). The reduced life expectancy in MPN is largely driven by thromboembolic events, as well as by risk of disease progression and leukemic transformation (Tefferi et al. 2014; Girodon et al. 2010; Wolanskyj et al. 2006; Hultcrantz et al. 2015; Leontyeva et al. 2026).

Previous studies assessing plasma proteins in MPN disease and MPN subtypes have largely focused on selected inflammatory mediators, most often using candidate-based low-throughput assays such as ELISA or multiplex cytokine panels (Mambet et al. 2018; Tefferi et al. 2011; Baldauf et al. 2025; Rai et al. 2022; Rahman et al. 2022). For example, Increased levels of IL-6 in MPN patients have been widely reported (Bourantas et al. 1999; Hsu et al. 1999; Panteli et al. 2005; Boissinot et al. 2011; Vaidya et al. 2012; Pourcelot et al. 2014; Cacemiro et al. 2018); Tefferi *et al*. reported increased plasma levels of IL-8, IL-2R, IL-12, and IL-15 in PMF compared to healthy controls (Tefferi et al. 2011), and Mambet *et al*. reported that Dickkopf-related protein 1 (DKK1) levels were increased significantly in PV and PMF compared to ET (Mambet et al. 2018). Furthermore, High mobility group box 1 (HMGB1) has been reported to be elevated in both serum and bone marrow mononuclear cells from MPN patients compared to patients with iron deficiency anemia (Wu et al. 2025).

The clinical and biological heterogeneity of MPN is partly shaped by the underlying driver mutation, and this has also been reflected at the blood protein levels, as increased CXCL10/IP-10 (Interferon gamma-induced protein 10, IP-10) levels have been linked to *JAK2* mutation in PMF(Cacemiro et al. 2018), and apolipoprotein A1 (APOA1) levels have been reported to be associated with *JAK2* mutation and correlated with the mutation burden in PV (Mossuz et al. 2007). In addition, CALR protein level has been shown to be overexpressed in *JAK2* mutated granulocytes, indicating that driver mutations may influence broader proteomic programs beyond the mutated gene itself (Socoro-Yuste et al. 2017).

More recently, large-scale population studies have shown increasing evidence for a role of plasma proteins in the risk for developing MPN. Xiong *et al*. demonstrated that elevated Interleukin-2 receptor alpha subunit (IL2RA) and CXCL10 levels were associated with increased MPN risk (Xiong et al. 2024), and, using data by the UK Biobank, Tran *et al*. identified 115 plasma proteins significantly associated with the future risk of developing myeloid neoplasms (Tran et al. 2024). However, broader high-throughput proteomic studies remain limited, and the extent to which plasma protein profiles reflect MPN- intrinsic biology and can provide insights into disease biology and subtype biomarkers, is still incompletely understood.

In addition, all of the aforementioned studies are limited by the use of healthy controls with normal blood counts. Eriksson *et al*. identified significant associations between platelet count / white blood cell count / erythrocyte volume fraction and cytokines (Eriksson et al. 2025), clearly demonstrating that blood cell counts *per se* alter the plasma proteome. To assess the inherent plasma proteome at diagnosis of MPN disease, we chose non-MPN controls consisting of individuals referred to the Department of Hematology due to suspicion of MPN who did not fulfil the diagnostic criteria. This design minimized confounding from elevated blood counts alone and facilitated the identification of plasma protein signatures more specifically associated with MPN biology.

## Materials and methods

### Patient cohort

We included consecutive adult patients referred to the Department of Hematology due to suspected diagnosis of MPN, from March 2022 to February 2025. Exclusion criteria were ongoing cytoreductive treatment, and inability to give informed consent. Peripheral blood samples were collected at the first outpatient clinic visit (baseline) for all patients.

This study was approved by the regional ethical review board in Stockholm, Sweden (reference number 2019-05192, 2022-01828-02) and conducted in accordance with the Declaration of Helsinki. All participants provided oral and written informed consent.

### Laboratory parameters and diagnostic information

We collected information from medical charts on comorbidities including previous thromboses, where thrombosis was defined as any type of arterial or venous thrombosis before the referral to the Department of Hematology. Results from molecular testing at MPN diagnosis were also collected, including the allelic burden of *JAK2*, *CALR*, and *MPL* mutations assessed by next-generation sequencing. In five patients, a *JAK2* mutation burden was reported as <5% but confirmed by RT-PCR with a cut-off of ≥0.5% for confirmed presence of the *JAK2* V617F mutation; for LASSO regression analysis, these values were set to 4%.

### Sample processing

Whole blood was collected in citrate tubes and centrifugated 10 minutes at room temperature, at 2000x g. Supernatant was then pooled and subsequently centrifugated again under the same conditions, whereafter the platelet poor plasma was aliquoted and stored at -80°C until thawed for proteomic profiling.

### Proteomics assay

Plasma proteomics profiling of blood samples was performed with Olink Explore 384 platform using Cardiometabolic (N=369) and Inflammation (N=368) panels, in total 737 unique proteins. Three proteins; IL-6, TNF and CXCL8, overlapped between these two panels. Relative protein concentration was expressed as NPX (log_2_ normalized protein expression). The Olink assay, data processing and quality control procedures were performed as previously described (Söderlund et al. 2023).

### Statistical analysis

To assess data distribution, we applied the Shapiro-Wilk test to examine whether the plasma protein NPX values followed a normal or log-normal distribution. Differential expression analyses between MPN and non-MPN controls were performed using the olink_ttest function implemented, and adjusted age and gender with olink_lmer function in the OlinkAnalyze package (Nevola et al. 2022). Multiple testing correction was conducted using the Benjamini&Hochberg (BH) method, and proteins with an adjusted p-value < 0.05 were considered significantly differentially expressed.

To further assess associations between plasma proteins and MPN subtypes (ET, PV, and PMF), we applied a Cox proportional hazards model with a least absolute shrinkage and selection operator (LASSO) penalty to compare MPN overall, ET, PV and PMF to non-MPN controls, respectively. Covariates included age, sex, and blood count parameters (platelet, leukocyte, and hemoglobin levels).

The significance of each protein feature was determined by its median coefficient across 1000 bootstrap replicates of the LASSO Cox regression. Proteins that retained a nonzero coefficient in ≥700 out of 1000 replicates were considered robustly associated with MPN. These proteins were further evaluated using the Kruskal–Wallis test and BH-adjusted p-values to determine their statistical significance within MPN and its subtypes.

Receiver operating characteristic (ROC) curve analysis was used to evaluate the performance of plasma EPO, TFRC and their combination for predicting PV among MPN patients. Patients were classified as PV or non-PV based on clinical diagnosis. ROC curves were generated separately for EPO and TFRC, and a combined model including both EPO and TFRC was constructed using logistic regression. The predicted probabilities from the combined model were then used for ROC analysis. The area under the curve (AUC) was calculated for each model to assess diagnostic performance.

### Software and Algorithms

All statistical analyses and data visualization were performed using R (version 4.5.0). R package OlinkAnalyze (v 4.3.1) was used for the analysis of proteomic data from Olink.

### Data and code availability

The plasma proteomics data generated in this study will be deposited in the SciLifeLab Data Repository upon publication. Due to Swedish Law and local regulations on sensitive personal information, the data is available under restricted access. R codes used to analyse and generate figures in this study are posited in Github (https://github.com/joey0214/2026_MPN_plasma_proteomics).

## Results

### Patient characterization

The scheme of the study design is illustrated in Figure 1 A. Of the 87 individuals included, 55 received an MPN diagnosis, comprising 30 ET, 16 PV and 9 PMF cases, while 32 did not fulfill diagnostic criteria and were included as non-MPN controls (Figure 1A).

**Figure 1.**
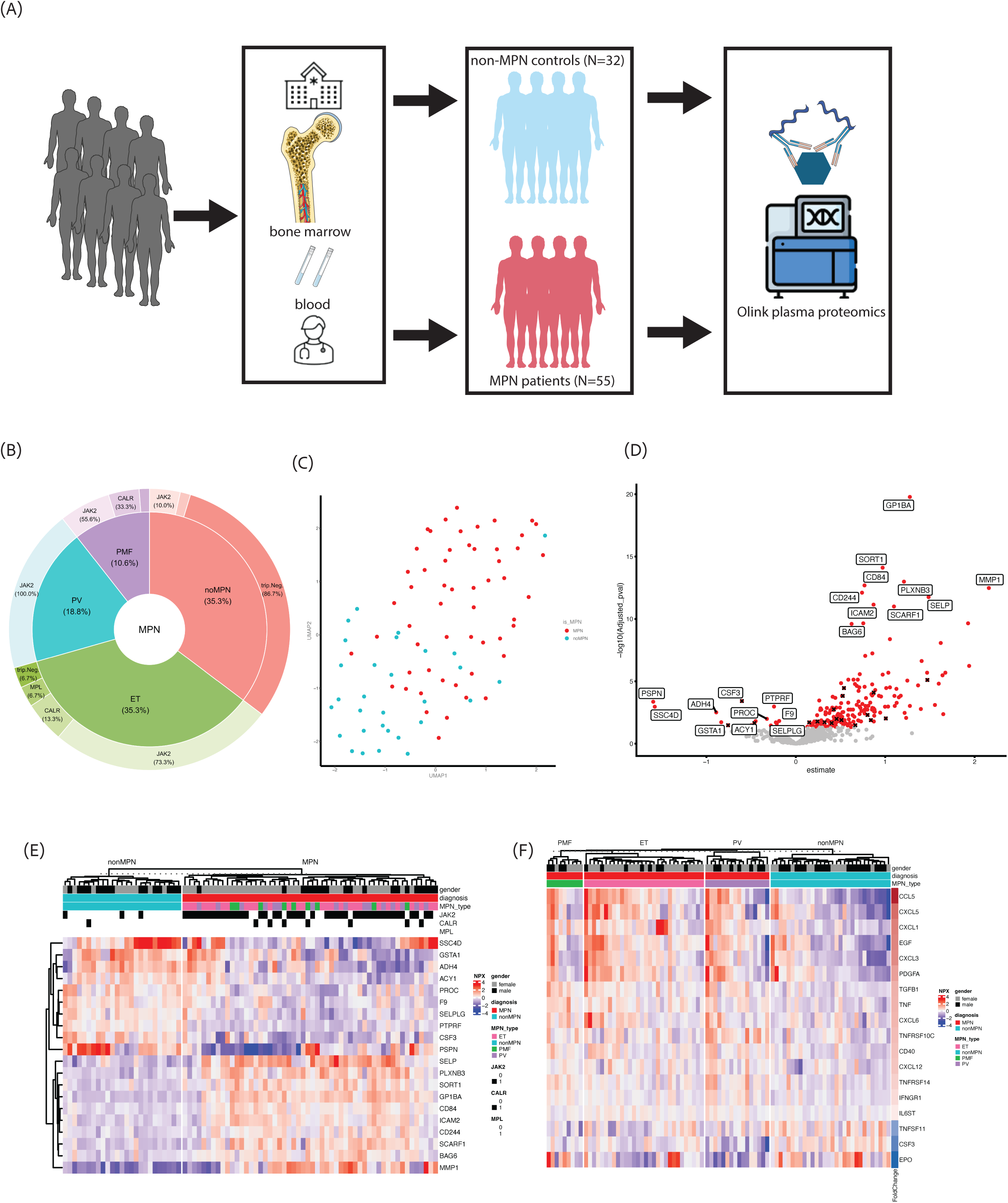
(**A**) Scheme overview of the study design. (**B**) Pie chart mutation profiles of included individuals. (**C**) UMAP of all MPN and non-MPN samples across all time points, with points colored by sample type. Red dots for MPN patients; blue dots for non-MPN controls. (**D**) Volcano plot of differential plasma protein expression in MPN versus non- MPN controls. The top 10 (most significance by adjusted p-value) up- and downregulated proteins are labeled with protein name. Dot color represents adjusted p-value significance (red for adjusted p-value < 0.05, gray for adjusted p-value > 0.05). In total, 189 proteins were significantly different between groups after age and gender adjustment: 16 proteins (9 Cardiometabolic, 7 Inflammation) were lower in MPN, and 173 proteins (88 Cardiometabolic, 85 Inflammation) were higher in MPN. 18 significant differential expressed cytokine proteins were marked with black crosses. (**E**) Heatmap showing the top up- and downregulated proteins in MPN vs non-MPN controls. Columns are annotated with gender, MPN diagnosis, MPN subtype and mutation profiles. (F) Heatmap showing the significant differential expressed cytokine proteins in MPN vs non-MPN controls. Columns are annotated with MPN diagnosis and MPN subtype. Rows are annotated with FoldChange from the comparison between MPN and non-MPN patients.

Patient characteristics were summarized in Table 1. Among the 55 MPN patients, 36 (65%) were female, whereas 11 (34%) of the 32 non-MPN controls were female. This gender imbalance was primarily driven by the ET subtype, in which 22 (73%) of the 30 ET patients were female (Supplementary Figure 1 A). A similar proportion of individuals in each group had a history of thrombosis (whether it was arterial or venous): 16 (29%) of 55 MPN patients and 7 (22%) of 32 non-MPN controls (Supplementary Figure 1 B).

**Table 1.**
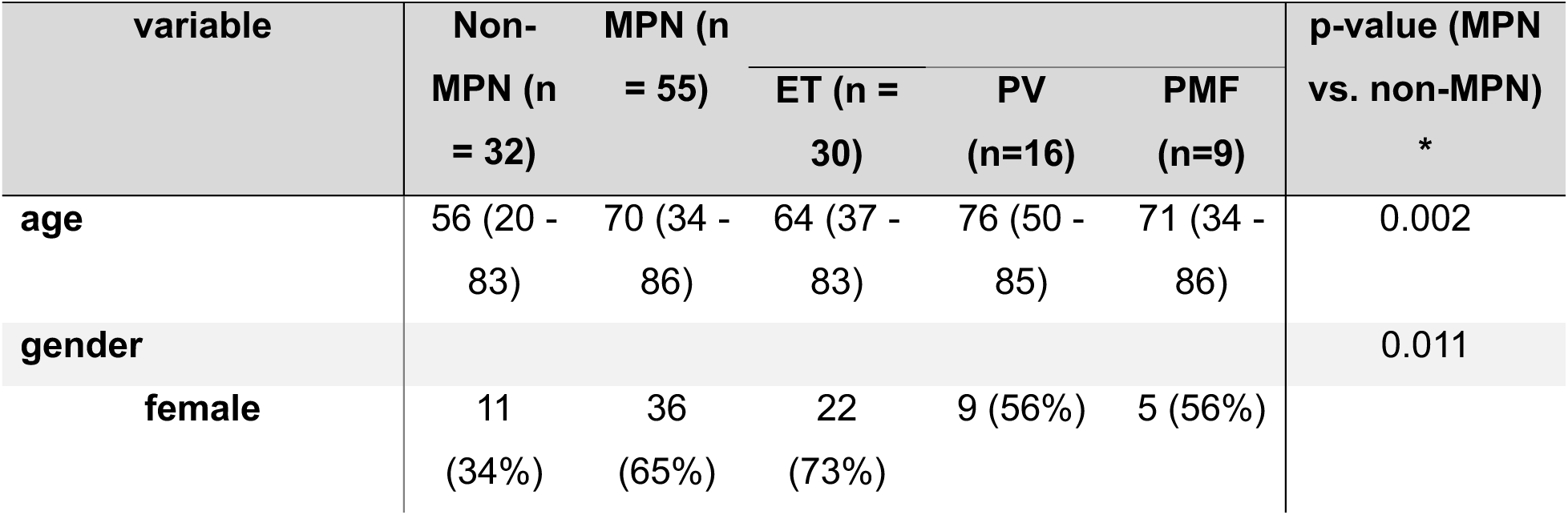
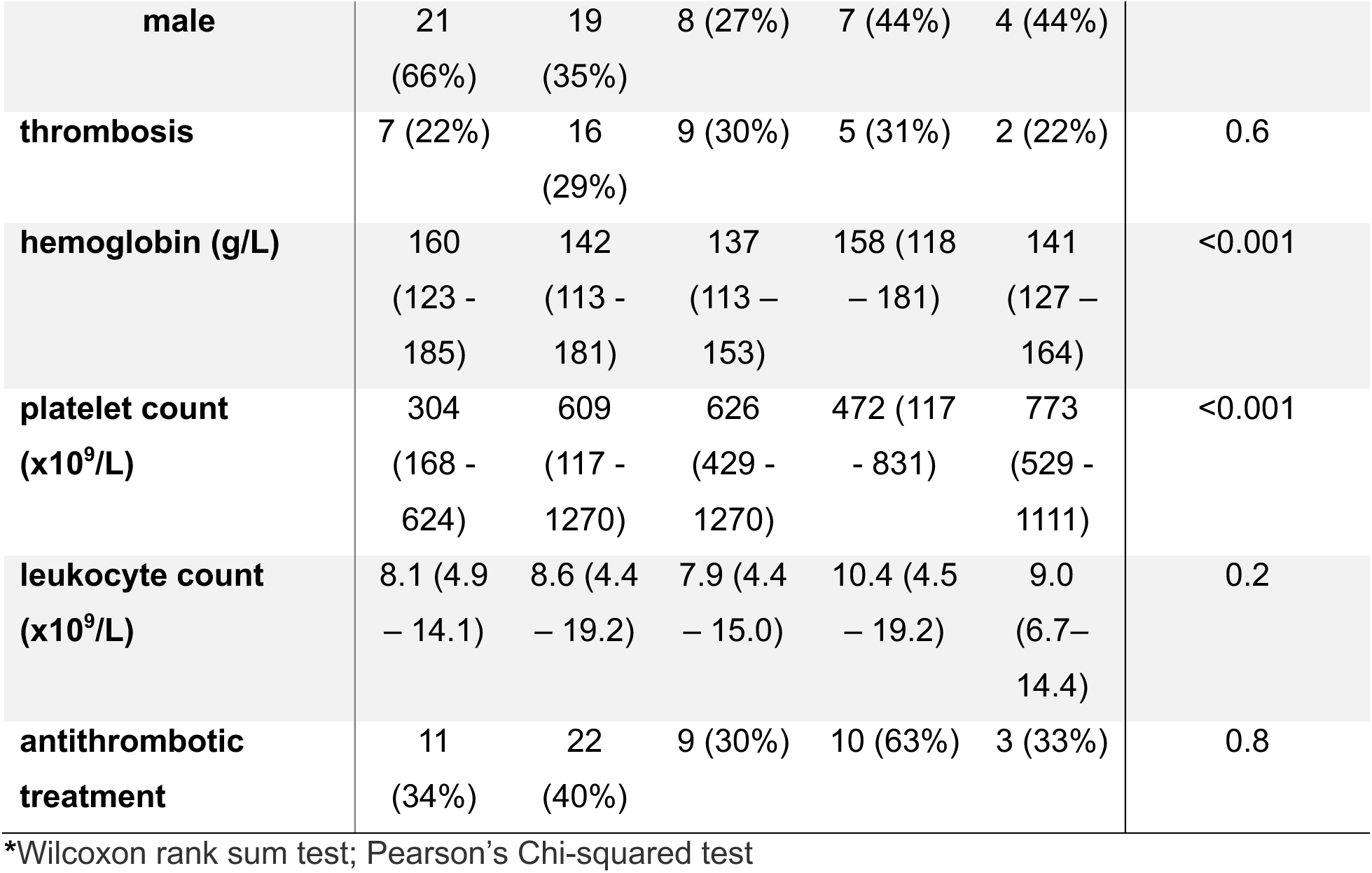
Baseline characteristics of the cohort.

Among the 55 MPN patients, *JAK2* mutation was found in 43 (78%), *CALR* in 7 (13%) and *MPL* in 3 (5.5%) cases. In the non-MPN group, *CALR* mutation was found in 1 and *JAK2* in 3 cases, these cases could be classified as CHIP (Figure 1B, Supplementary Table 1). The proportion of patients who received antithrombotic treatment was comparable between the two groups.

Hemoglobin and platelet counts differed significantly between non-MPN controls and MPN patients but were relatively homogeneous within the two groups (Supplementary Figure 1 C-E). PV patients had hemoglobin levels similar to non-MPN, while ET and PMF patients exhibited lower values. PV cases exhibited higher leukocyte counts than the other MPN subtypes and controls. ET and PMF cases had higher platelet counts than PV patients and non-MPN controls.

### Plasma proteins elevated in MPN patients compared with non-MPN controls

Two non-MPN controls failed data quality control according to Olink data handling (Supplementary Figure 2 A-D) and were excluded from further analysis.

Three proteins, TNF, IL-6 and CXCL8, were included in both the Cardiometabolic and the Inflammation panels and correlated well between panels (Supplementary Figure 2 E-G). Therefore, the NPX values from the Cardiometabolic panel were used for downstream analysis. The UMAP visualization demonstrates clear separation between MPN and non-MPN controls, indicating distinct plasma proteomic profiles between the groups (Figure 1 C).

After correcting for age and sex bias between the MPN and non-MPN groups using linear mixed-effects model (Supplementary Figure 3 A), differential expression analysis identified 189 proteins significantly (adjusted p-value < 0.05) altered between MPN and non-MPN controls (Figure 1 D, Supplementary Table 2). Of the 189 differentially expressed plasma proteins, 16 were downregulated in MPN whereas 173 proteins were upregulated. The NPX levels of the top 10 (most significant by adjusted p-value) up- and down-regulated proteins in Figure 1 D were shown in the heatmap Figure 1 E, which includes column annotations for gender, MPN status and MPN subtype.

Interstitial collagenase (MMP1) was the most strongly up-regulated plasma protein (Figure 1 D). MMP1 levels were significantly different between MPN – including all subtypes – and non-MPN controls, while no significant difference was observed among the MPN subtypes themselves (Supplementary Figure 3 B). Glycoprotein Ib platelet subunit alpha (GP1BA) was the most statistically significant up-regulated plasma protein (Figure 1 D) and has previously been associated with MPN risk in the UK BioBank study (Tran et al. 2024). GP1BA was significantly elevated in MPN patients compared with non-MPN controls. Within the MPN subtypes, differences were observed only between ET and PMF (Supplementary Figure 3 C). Overall, the plasma proteomic profiles demonstrated clear differences between MPN and non-MPN controls, as well as heterogeneity across MPN subtypes.

### Altered cytokines between MPN and non-MPN patients

Of the 189 differentially expressed proteins, 18 proteins were annotated to the KEGG cytokine and cytokine receptor interaction pathway (hsa04060) and were indicated by red dots with black crosses in Figure 1 D, including CCL5, CD40, CSF3, CXCL1, CXCL12, CXCL3, CXCL5, CXCL6, EGF, EPO, IFNGR1, IL6ST, PDGFA, TGFB1, TNF, TNFRSF10C, TNFRSF14 and TNFSF11. The expression patterns of these proteins were shown in the heatmap in Figure 1 F. Notably, cytokines related proteins previously implicated in MPN, including TNF, TGFB1, were identified among the differently expressed proteins between MPN and non-MPN controls, whereas IL-6 was not significantly altered.

Notably, although there was some degree of overlap in EPO levels, EPO was significantly lower in PV compared with non-MPN controls (Supplementary Figure 3 D). The EPO level showed variation between MPN subtypes as well (Supplementary Figure 3 E).

In the STRING protein-protein interaction plot for the 18 cytokine related proteins that were significantly differentially expressed between MPN and non-MPN patients, TNF had a central role (Supplementary Figure 3 F). Among these 18 cytokine-related proteins, CCL5, CXCL1, and CXCL5 showed the most pronounced alterations and exhibited the highest number of interactions within the network.

### Functional pathway enrichment reveals upregulation of hemostasis and platelet-related proteins in MPN plasma

To investigate the biological processes associated with dysregulated plasma proteins, we performed functional overrepresentation analysis. Because the study used targeted Olink protein panels with a predefined set of measurable proteins, we applied an approach that accounts for panel-specific background gene sets in the hypergeometric test. The analysis revealed significant enrichment of the Reactome pathways “hemostasis” and “platelet activation, signaling and aggregation” among the differential expressed proteins (Supplementary Figure 4 A). To validate these findings, we examined NPX values for all proteins annotated to these pathways. Proteins involved in the “hemostasis” pathway showed a clear difference in expression between MPN subtypes and non-MPN controls, with modest variability also observed among MPN subtypes (Supplementary Figure 4 B). Proteins associated with “platelet activation, signaling and aggregation” similarly differed between MPN subtypes and non-MPN controls, but no difference was detected within MPN subtypes (Supplementary Figure 4 C).

The overall NPX levels of proteins from KEGG cytokine and cytokine receptor interaction pathway showed differences between MPN and non-MPN (Supplementary Figure 4 D), as well as between subtypes (Supplementary Figure 4 E). However, no cytokine related pathways were found to be significant in functional enrichment analysis.

### Plasma protein alterations associated with *JAK2* mutation

To assess whether the *JAK2* mutation was driving specific plasma protein patterns, and associated with cytokine profiles, we performed the protein differential expression analysis comparing *JAK2* mutant MPN to *JAK2* wildtype MPN. We found a significant upregulation of IL-6 and GH1 in *JAK2* mutant MPN (Figure 2 A). The IL-6 was clearly driven by *JAK2* mutation and not by MPN disease subtype (Figure 2 B-C) and a similar pattern was found with GH1 (Supplementary Figure 5 A-C). The up-regulated trend of IL- 6 and GH1 (Figure 2 D) was also seen, though not significant, when comparing *JAK2*- mutant ET (N=22) to *JAK2* wildtype ET (n=8). When comparing *JAK2* mutant ET to *JAK2* mutant PV, EGFR and GDF2 were significantly increased in ET and TFRC significantly increased in PV (Figure 2 E). Furthermore, we examined the associations between driver mutation status and peripheral blood counts at diagnosis. Driver mutations showed significant associations with platelet counts, while gender was associated with hemoglobin levels (Supplementary Figure 5 D). These findings support that mutational background contributes to variation in blood count parameters among MPN patients. Together, these findings suggest that plasma protein profiles are not strongly influenced by *JAK2* mutation status.

**Figure 2.**
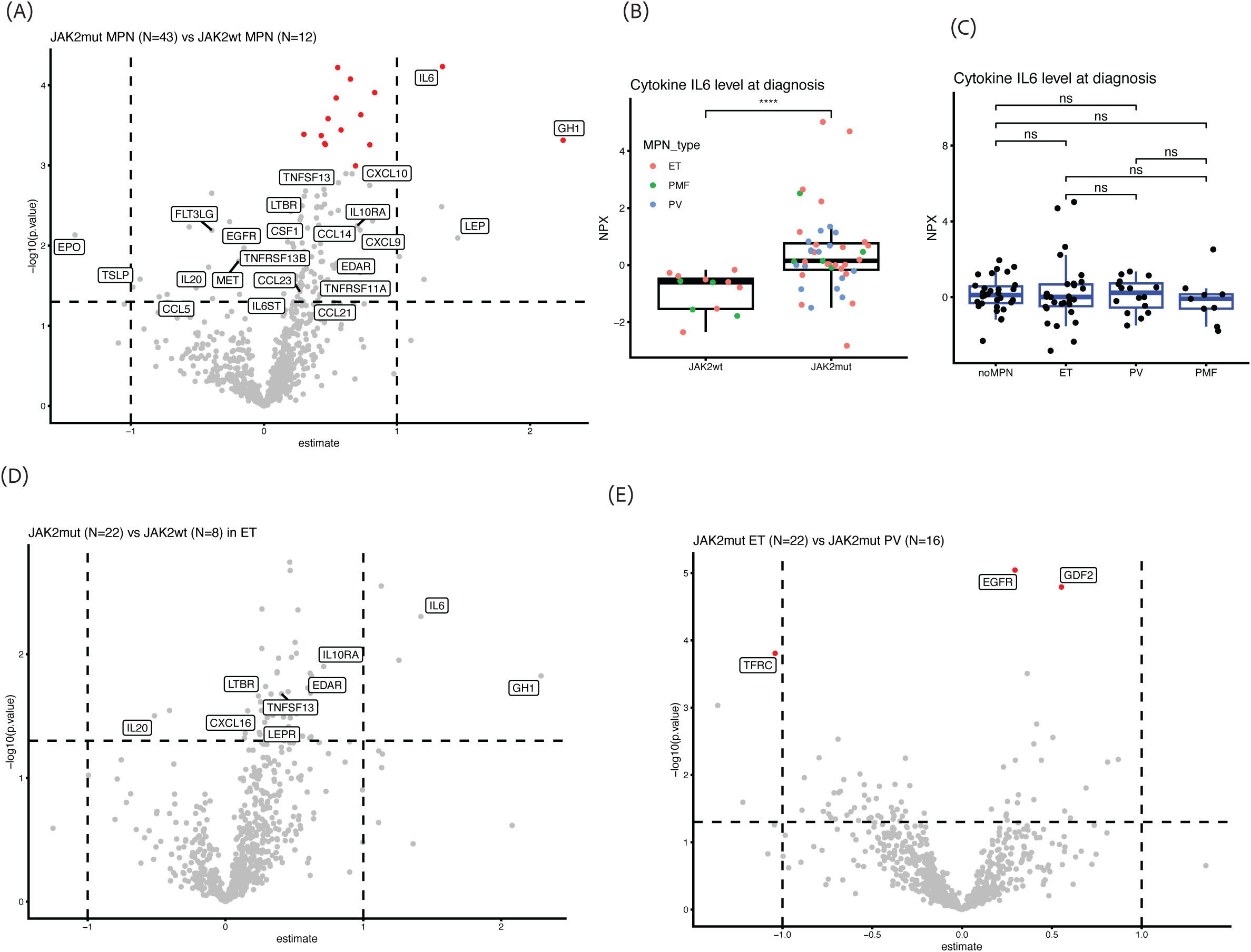
(**A**) Volcano plot showing differential protein expression between *JAK2*-mutant (N=43) MPN and *JAK2*-wildtype MPN (N=12). IL-6 and GH1 were the only significant differentially expressed cytokine proteins. Boxplots of IL-6 plasma level in *JAK2*-mutant and wildtype MPN patients (**B**); as well as in MPN subtypes (**C**). Pairwise comparisons were performed using t-test. Significance levels: ns: p > 0.05; *: p <= 0.05; **: p <= 0.01; ***: p <= 0.001; ****: p <= 0.0001. (**D**) Volcano plot showing differential protein expression between *JAK2*-mutant ET (N=22) and *JAK2*-wildtype ET (N=8). (**E**) Volcano plot showing differential protein expression between *JAK2*-mutant ET (N=22) and *JAK2*- mutant PV (N=16). Transferrin receptor (TFRC), epidermal growth factor receptor (EGFR) and Growth differentiation factor 2 (GDF2) were significantly differentially expressed.

### Plasma proteins associated with MPN and MPN subtypes

We applied Cox LASSO regression to evaluate associations between plasma protein and MPN status, comparing non-MPN to MPN, ET, PV and PMF, respectively (Figure 3 A). Overall, the results suggest that MPN-associated proteins were both partly shared among the subtypes but were also subtype-specific. Five proteins were significant predictors of MPN overall, among which GP1BA and sortilin (SORT1) were the two strongest, supporting their role as general MPN biomarkers. SORT1 was associated with both ET and PV as well. GP1BA was not only differentially expressed between MPN and non-MPN controls (Figure 1 D), but also significantly altered across MPN subtypes compared with non-MPN controls (Supplementary Figure 3 C). PV displayed the most distinct proteomic profile, including increased levels of TFRC, SORT1 and semaphorin 7A (SEMA7A) (Figure 3A-D). Notably, GDF2 was associated with PV differently, with negative coefficients (Figure 3A, E), suggesting lower levels or inverse association with PV. In contrast, ET and PMF showed fewer subtype-specific markers, with GP1BA strongly associated with both ET and PMF, consistent with platelet- and hemostasis-related biology. PMF is mainly characterized by GP1BA and SCARF1, suggesting a more limited but still distinct protein signature.

**Figure 3.**
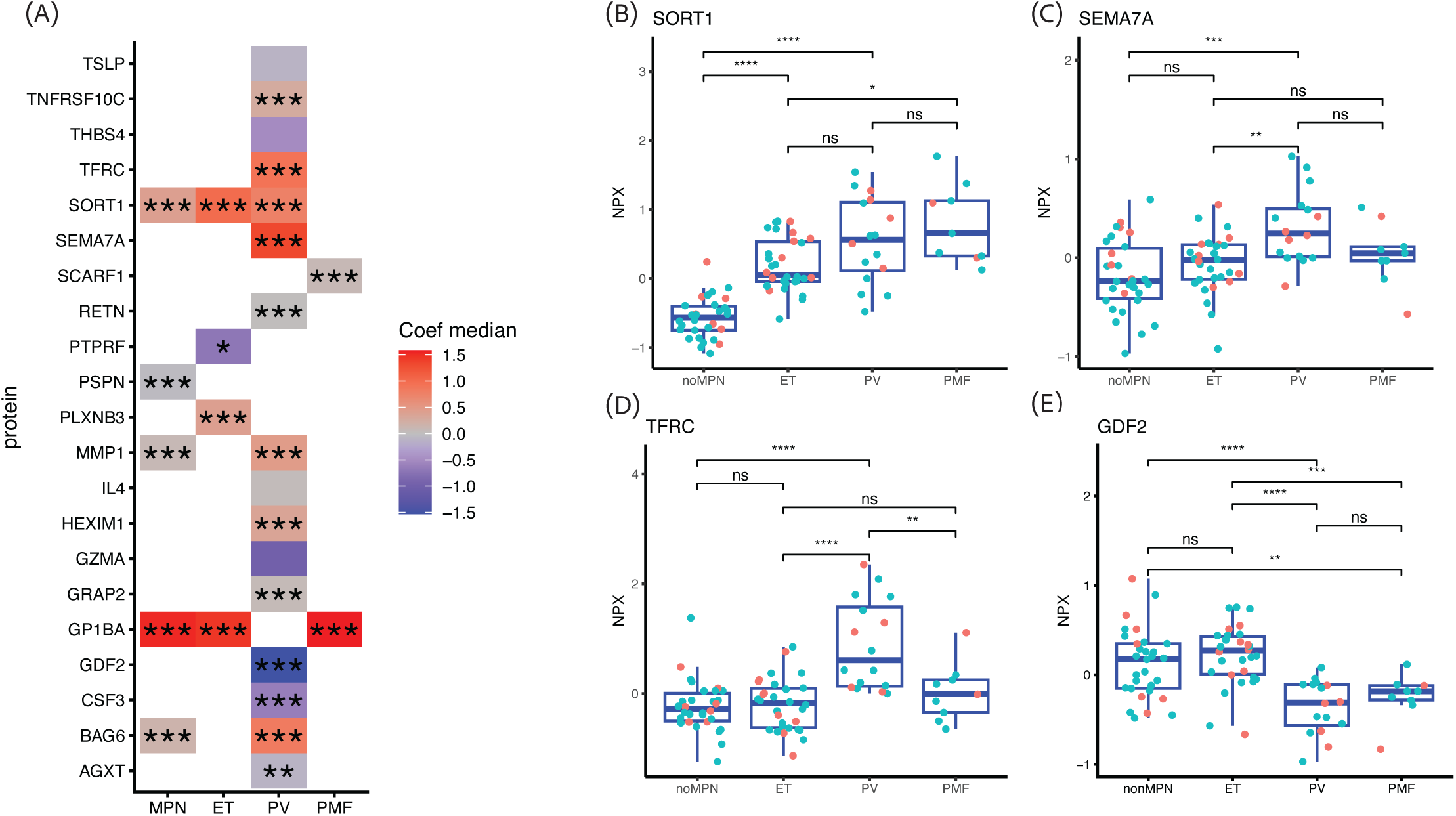
MPN subtype specific associated plasma proteins. (**A**) The association between plasma proteins and MPN/subtypes assed by binomial LASSO regression. Significance levels from Kruskal-Wallis test with Benjamini-Hochberg adjusted p-value: ns: adj.p > 0.05; *: adj.p <= 0.05; **: adj.p <= 0.01; ***: adj.p <= 0.001; ****: adj.p <= 0.0001. (**B**-**E**) Boxplots of sortilin (SORT1), Semaphorin-7A (SEMA7A), Transferrin receptor protein 1 (TFRC) and GDF2, expression in plasma at diagnosis. Pairwise comparisons were performed using t-test. Significance levels: ns: p > 0.05; *: p <= 0.05; **: p <= 0.01; ***: p <= 0.001; ****: p <= 0.0001.

With these associations between plasma proteins and MPN, we further investigated their performance of predicting MPN and MPN subtypes. Firstly, a significant correlation between GP1BA level in plasma and platelet count was found in ET (r2=0.31, p- value=0.0015) and in PMF (r2=0.62, p-value=0.011) (Supplementary Figure 6 A). SEM17A was correlated with leukocyte count in non-MPN patients (r2= 0.3, p-value= 0.0016) and PV (r2= 0.35, p-value= 0.016) (Supplementary Figure 6 B). TFRC expression did not correlate with hemoglobin, leukocyte count or platelet count (Supplementary Figure 6 C). Therefore, TFRC was the candidate for further investigation.

With the lower level of EPO in PV (Supplementary Figure 3 D-E, Supplementary Figure 6 D), combining EPO and TFRC together was a very good discriminator of PV from the other subtypes of MPN and non-MPN (Figure 4 A), without any observed correlation between EPO and TFRC level in plasma (Supplementary Figure 6 E). Receiver operating characteristic (ROC) curve analysis was performed to evaluate the performance of plasma EPO and TFRC protein levels to distinguish patients with PV from others (non-MPN, ET and PMF). The area under the curve (AUC) was calculated for EPO alone, TFRC alone, and combined EPO and TFRC, to assess diagnostic performance. In this analysis, EPO showed an AUC of 0.774, TFRC showed an AUC of 0.893, and the combined TFRC + EPO model showed the highest performance with an AUC of 0.933.

**Figure 4.**
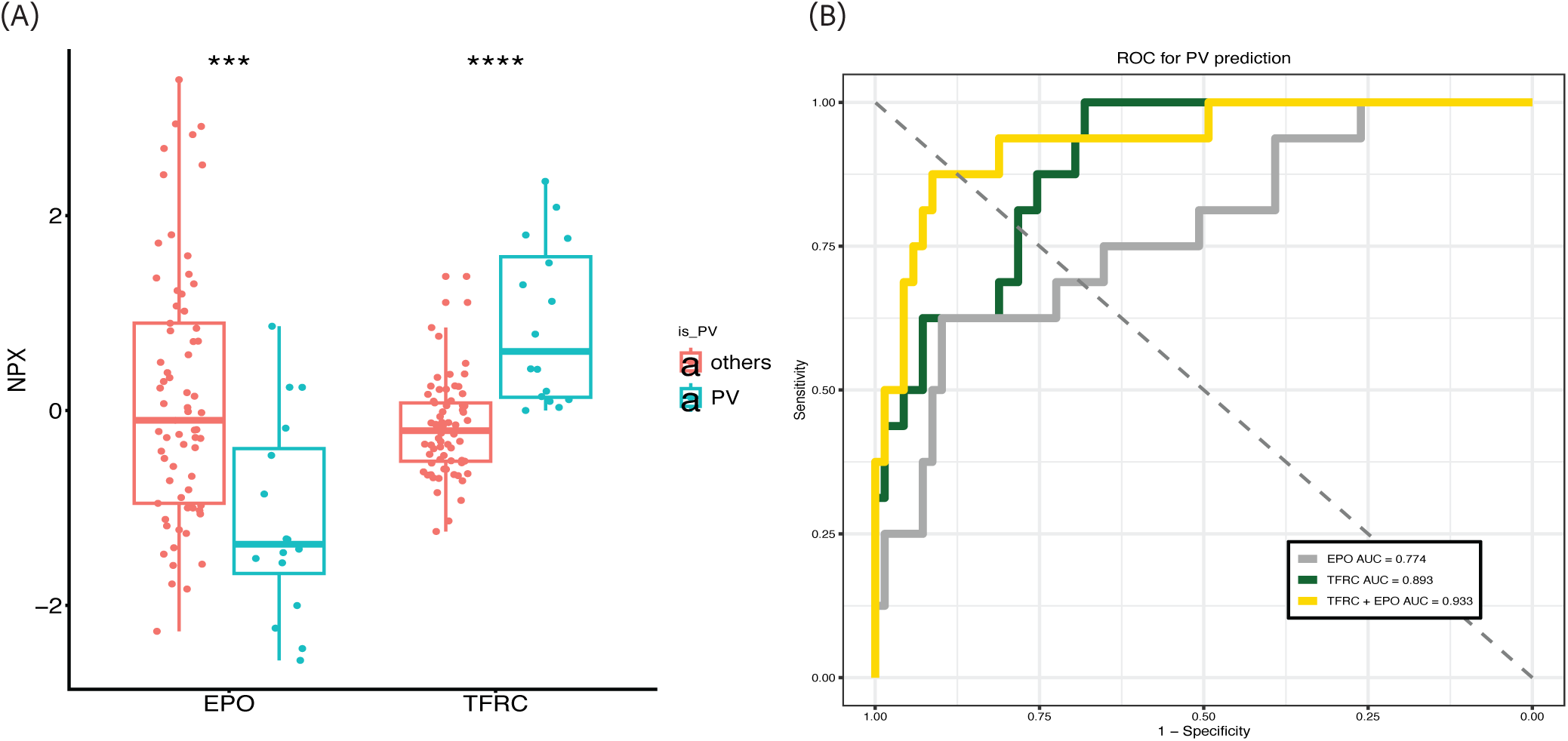
(**A**) The association between plasma proteins and MPN/subtypes assed by multinomial LASSO regression. The color represents the median coefficient: red indicates a positive association with the group, blue indicates a negative association, and asterisks indicate the significance of the association. Significance levels from Kruskal-Wallis test with Benjamini-Hochberg adjusted p-value: ns: adj.p > 0.05; *: adj.p <= 0.05; **: adj.p <= 0.01; ***: adj.p <= 0.001; ****: adj.p <= 0.0001. (**B**) Box plot showing plasma level of EPO and TFRC in MPN patients across at diagnosis. Red dots for EPO, and blue dots for TFRC. Pairwise comparisons were performed using t-test. Significance levels: ns: p > 0.05; *: p <= 0.05; **: p <= 0.01; ***: p <= 0.001; ****: p <= 0.0001. (**C**) ROV plot in PV with EPO (blue), TFRC (red) and combination of EPO and TFRC (green).

Among the 115 plasma proteins reported by Tran *et al*. to be associated with future MPN risk, where MPN developed on average seven years after blood sampling (Tran et al. 2024), 50 proteins were from the Olink Cardiometabolic and Inflammation panels used in our current study and thus could be compared between the cohorts. We found that of the 50 overlapping proteins, 26 proteins were significantly elevated in MPN compared to non-MPN at diagnosis in our current study (Figure 5 A), indicating that these proteins may be very early markers of MPN, elevated already years before diagnosis.

**Figure 5.**
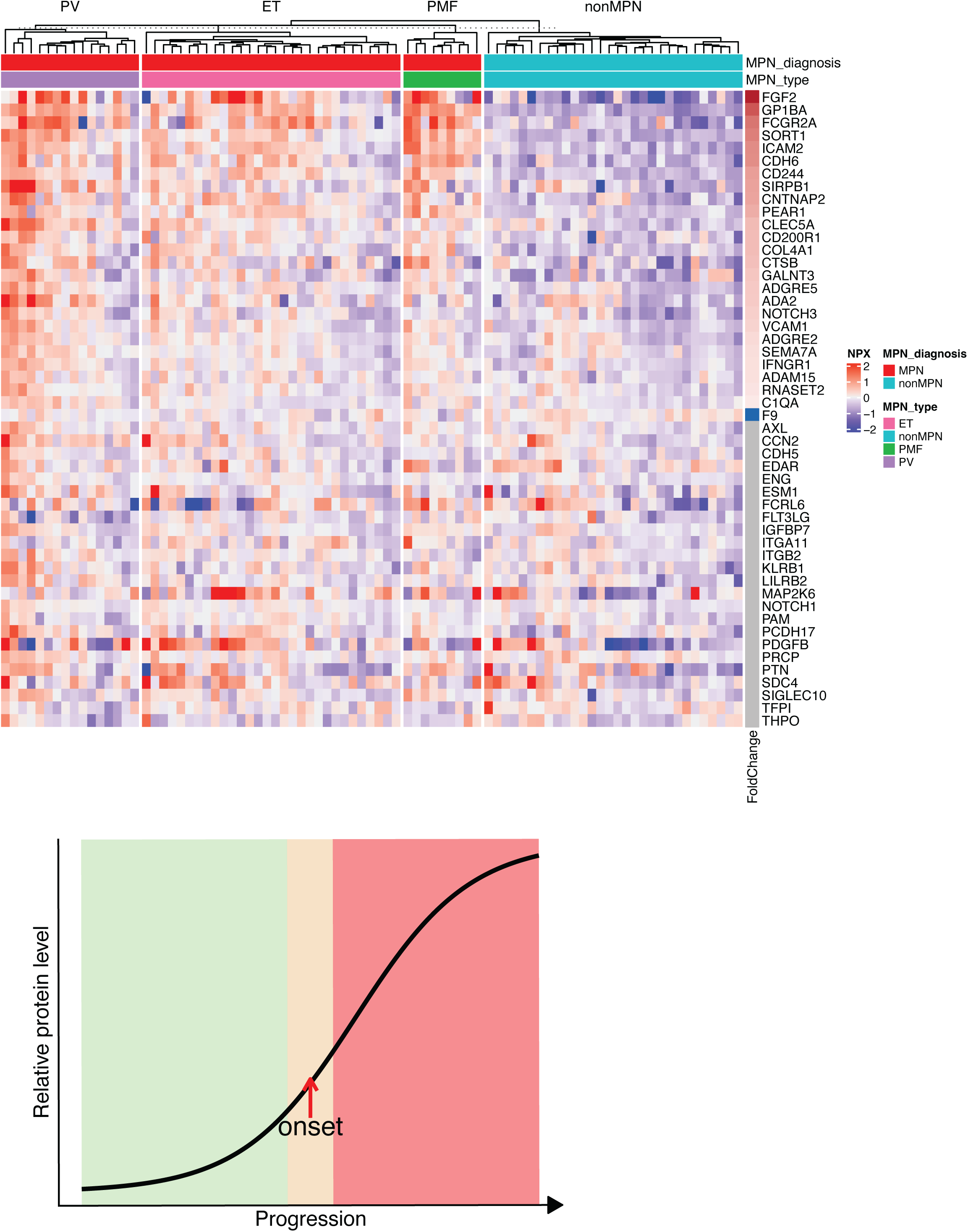
(**A**) Heatmap showing the expression level of 50 MPN-associated proteins in our Olink proteomic data from MPN and non-MPN patients. Columns are annotated with MPN diagnosis and MPN subtype. Rows are annotated with FoldChange from the comparison between MPN and non-MPN patients. (**B**) Schematic hypothesis showing a gradual increase in relative plasma protein levels during progression from a pre-diagnostic phase to clinically apparent disease onset. The green area indicates an early/preclinical phase with relatively low protein levels, the yellow area indicates the transition phase around disease onset, and the red area indicates established disease progression with markedly elevated protein levels. The red arrow marks the approximate point of clinical onset.

## Discussion

In this study, we demonstrated a distinct circulating protein signature in MPN, compared to non-MPN subjects with increased blood counts. After adjustment for age and gender, 189 plasma proteins remained differentially expressed, the majority being upregulated in MPN, and enriched for proteins in Hemostasis and Platelet activation, signaling, and aggregation pathways. This indicates that the plasma proteome at diagnosis reflects key biological features intrinsic to MPN, and may provide clinically relevant and specific information to assist the diagnostic process, compared to routine blood counts.

Eighteen of the 189 proteins were cytokines or cytokine-related proteins, of which several including TNF and TGFβ1 have been previously implicated in MPN pathobiology (Yao et al. 2022; Fleischman et al. 2011). However, the proinflammatory cytokine IL-6, that has been widely reported upregulated in MPN patients using ELISA and other low-throughput methods, was not significantly altered between MPN and non-MPN subjects in our study. Of note, studies finding increased IL-6 levels have mainly used healthy controls for comparisons, whereas no clear differences in IL-6 levels have been observed when comparing between MPN subtypes (Cacemiro et al. 2018; Pourcelot et al. 2014).

More recently, Eriksson *et al*. reported that higher IL-6 levels were strongly associated with platelet, white blood cell count, and neutrophil counts from 165 healthy individuals (Eriksson et al. 2025). Therefore, the increase in IL-6 demonstrated by previous studies may be associated with the increased blood counts *per se*, that are seen in MPN compared to healthy, and not an intrinsic feature of MPN disease. In the plasma samples of our cohort at diagnosis, IL-6 and GH1 were the two only cytokine-related proteins that significantly differed between *JAK2* V617F positive and negative MPN patients, indicating that IL-6 and GH1 were associated with the JAK2 V617F mutation. In contrast, previous studies showed no association between cytokine level and *JAK2* V617F mutation status (Cacemiro et al. 2018; Pourcelot et al. 2014).

Interestingly, Tran *et al*. identified 50 plasma proteins (of the Olink Cardiometabolic and Inflammation panels) predicting MPN risk (Tran et al. 2024), of which we could verify 26 as differentially expressed at the time of diagnosis in our MPN cohort. These were mainly proteins involved in platelet activation, monocyte and myeloid inflammation, and extracellular matrix remodeling. In contrast, proteins upregulated at diagnosis in our cohort, which were not associated with MPN risk in the study by Tran et al. and thus represent potential diagnostic markers, were primarily related to neutrophil activation, inflammasome signaling, and degranulation.

The LASSO regression analysis identified proteins with potential diagnostic and MPN subtype-specific relevance, where SORT1, GP1BA, PSPN, MMP1, and BAG6 distinguished MPN from non-MPN controls, while TFRC and SEMA7A were associated specifically with PV. These findings support a recent study demonstrating SORT1 to be upregulated in the platelet proteome of ET and PV patients using label-free LC-MS/MS (Kelliher et al. 2024), and GP1BA, SORT1 and SEMA7A were also found to be associated with MPN risk (Tran et al. 2024).

Our study provided additional MPN-subtype specific resolution, indicating TFRC as a specific biomarker of PV, independent of hemoglobin, platelet or white blood cell counts, suggesting that it captures biological information not reflected by routine blood parameters. TFRC has been described as a prognostic marker in multiple types of cancer, including in association with unfavorable outcome in cervical cancer (Wang et al. 2025), glioma (Schonberg et al. 2015), liver cancer (Hiromatsu et al. 2023), and renal cancer (Greene et al. 2017), and with favorable prognosis in lung cancer (Ren et al. 2025) in The Cancer Genome Atlas (TCGA) cohort.

Our results suggest that TFRC alone has a strong predictive power of PV, and when combining TFRC with serum erythropoietin, the predictive power for distinguishing PV from other MPN and non-MPN was 93.3%, indicating that this protein combination may provide a robust biomarker strategy for distinguishing PV from subjects with secondary erythrocytosis.

A major strength of our study is the use of a clinically relevant control group consisting of non-MPN individuals presenting with elevated blood counts and thus referred to the Department of Hematology on suspicion of MPN. By using this control group and not a healthy control group with normal blood counts, we remove the possible bias of detecting proteins associated with elevated blood counts *per se,* e.g. IL-6 that has been demonstrated to be associated with elevated white blood cell counts and platelet counts (Eriksson et al. 2025).

A limitation of our study is that although we assessed 737 unique proteins in a high-throughput assay, the number of proteins in the targeted Olink panels is restricted and may have excluded proteins potentially relevant for MPN. Future studies using deeper and broader protein coverage would be valuable for further elucidating disease mechanisms and improving biomarker discovery.

In conclusion, plasma proteomic profiling revealed a distinct circulating protein signature in MPN and identified distinct MPN-associated and subtype-specific signatures, with TFRC alone or in combination with serum erythropoietin showing potential as a blood biomarker of PV.

## Acknowledgements

Figure created with Servier Medical Art. We sincerely thank all participating patients and healthy volunteers for providing samples and contributing to this study.

## Contributions

J.U. and R.C. conceived the project. I.L. and A.R. collected and biobanked patient samples. I.L. conducted the chart review. X.Z. analyzed the data. X.Z. and J.U. wrote the manuscript. All authors reviewed and edited the manuscript. All authors read and approved of the final manuscript.

---

**Supplementary Figure 1.**
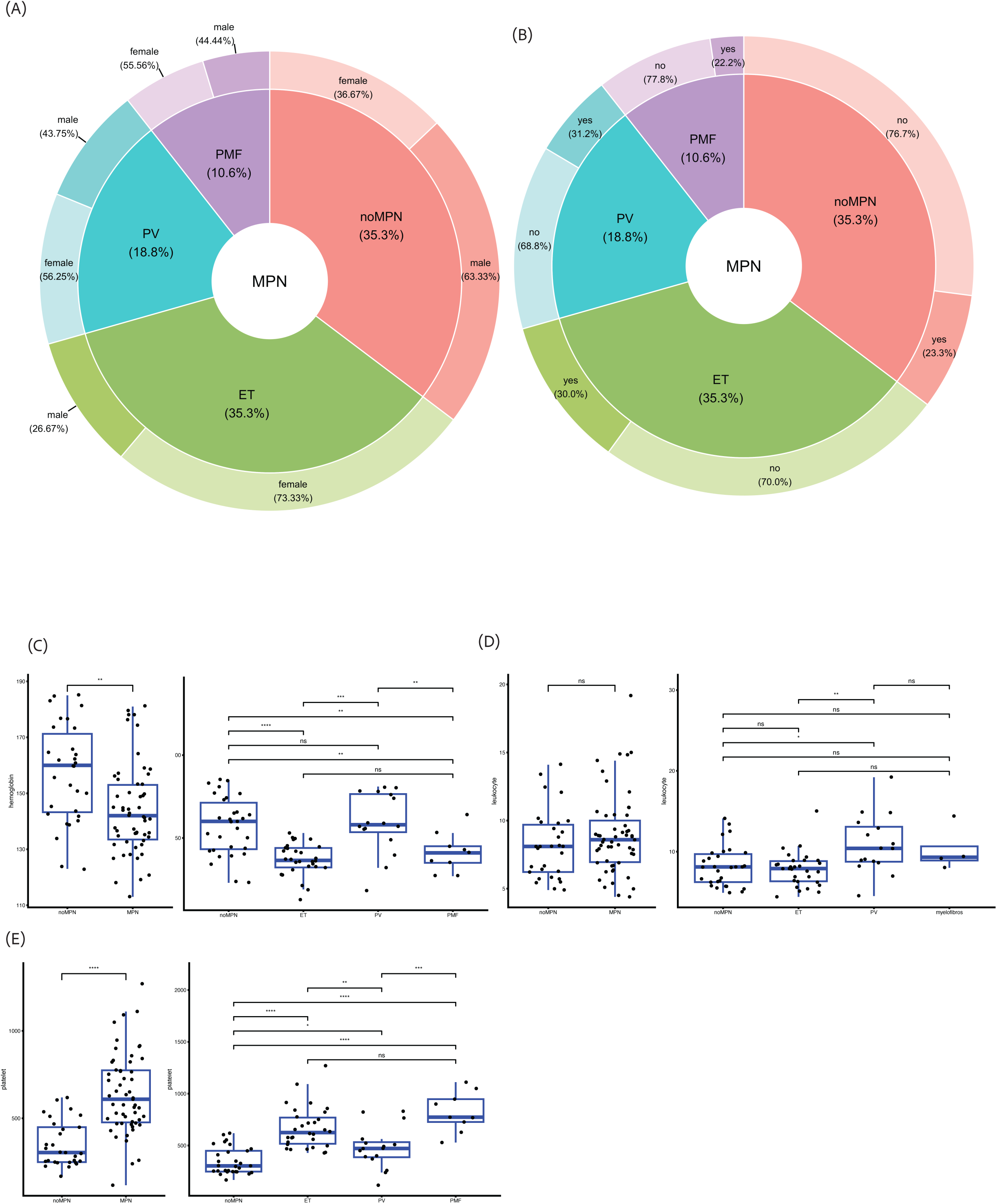
(**A**) Pie chart showing the distribution of samples by gender. (**B**) Pie chart showing the distribution of sample by thrombosis distribution. (**C-E**) The blood count of hemoglobin (**C**), leukocyte (**D**) and platelet (**E**) count in non-MPN controls and MPN samples at diagnosis, as well as their count in MPN subtypes.

**Supplementary Figure 2.**
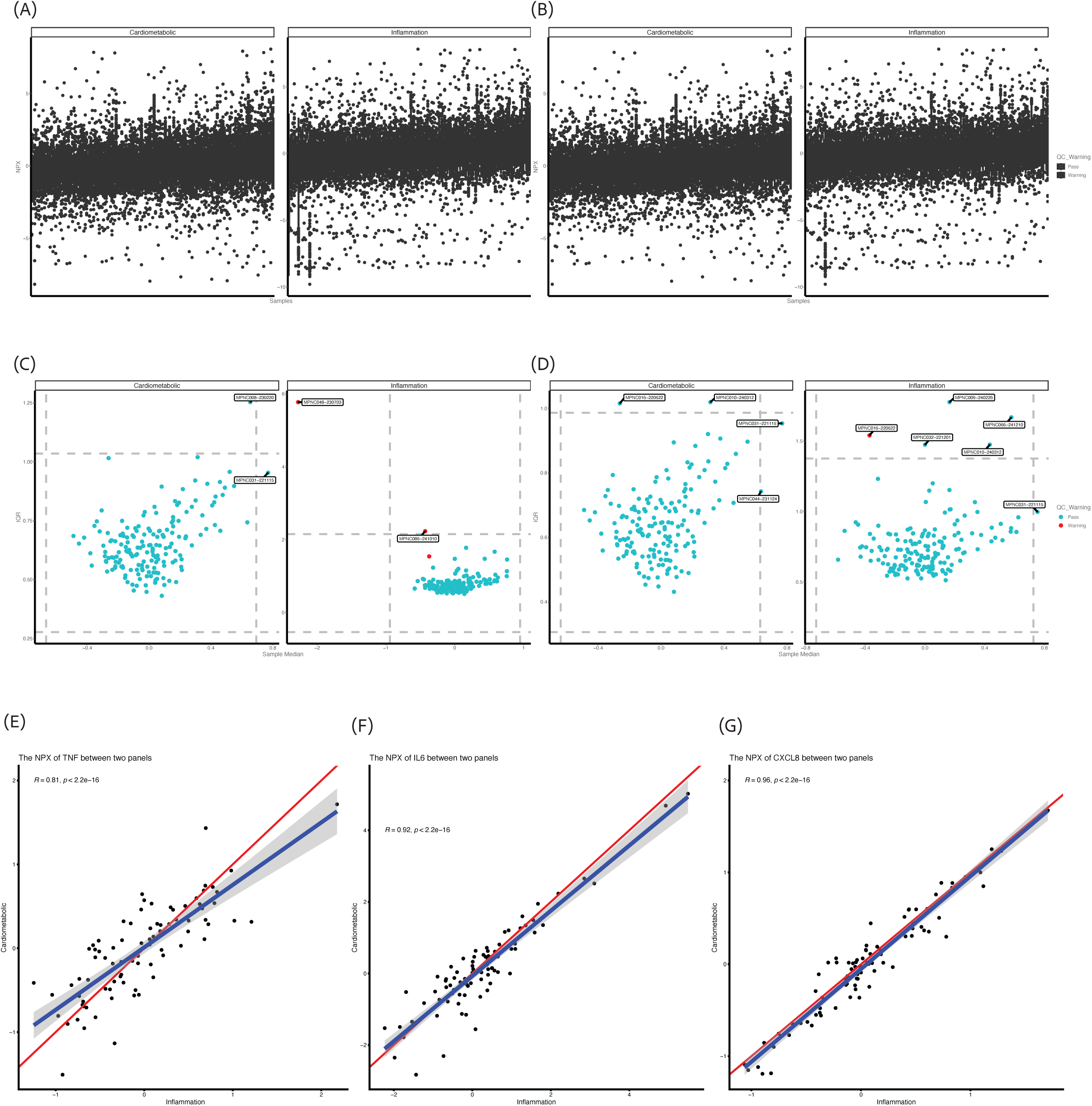
(**A**-**B**) NPX distribution per sample for the Cardiometabolic panel (**A**) and the Inflammation panel (**B**) before sample exclusion. (**C**-**D**) NPX distribution per sample for the Cardiometabolic panel (**C**) and the Inflammation panel (**D**) after sample exclusion. Linear correlation of plasma proteins TNF(**E**), IL-6 (**F**) and CXCL8 (**G**) between Cardiometabolic and Inflammation panel.

**Supplementary Figure 3.**
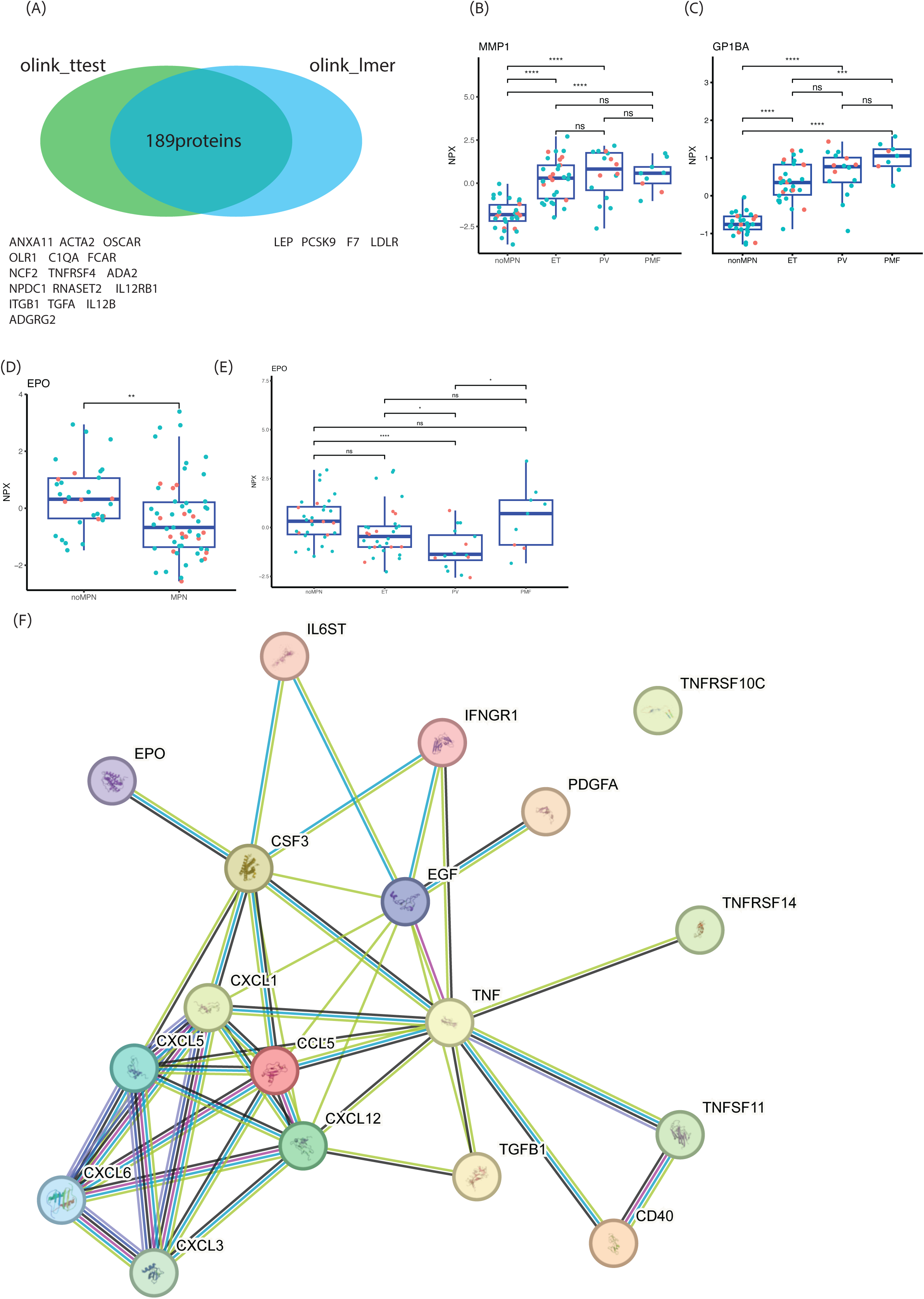
(**A**) Venn diagram of plasma protein differential expression analysis results using t-text and linear mixed model. (**B**) Box plot of MMP1 plasma level in non-MPN controls and MPN subtypes at diagnosis. (**C**) Box plot of GP1BA plasma level in non-MPN controls and MPN subtypes at diagnosis. Boxplots showing EPO plasma protein expression level in (**D**) non-MPN controls and MPN; (**E**) non-MPN controls and MPN subtypes. (**F**) Protein-protein interaction (PPI) network for these 18 cytokine related proteins was obtained from the STRING database. Nodes represent the proteins, and edges represent interaction between two proteins, edge colors represent evidence types. Only interactions with a high confidence (> 0.7) were considered.

**Supplementary Figure 4.**
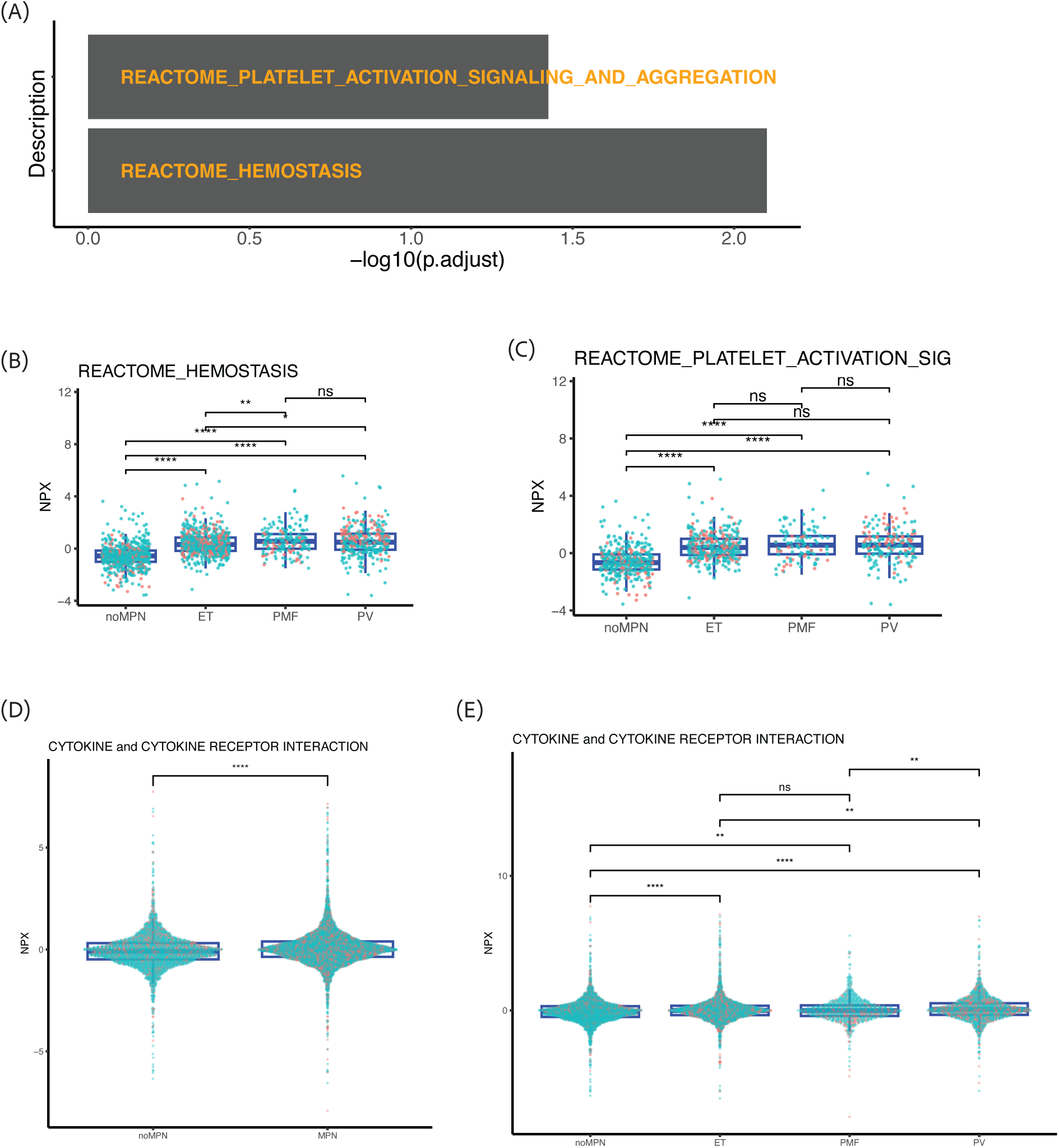
(**A**) Bar plot showing the two Reactome pathways significantly enriched in the Gene Set Enrichment Analysis (adjusted p-value < 0.05). (**B**-**C**) Boxplots of protein NPX levels for the pathways REACTOME HEMOSTASIS (**B**) and REACTOME platelet activation signaling and aggregation (**C**). Box plot of overall plasma level of cytokine and cytokine receptor interaction at diagnosis, between MPN and non-MPN controls (**D**), and between MPN subtypes (**E**). Significance levels: ns: p > 0.05; *: p <= 0.05; **: p <= 0.01; ***: p <= 0.001; ****: p <= 0.0001

**Supplementary Figure 5.**
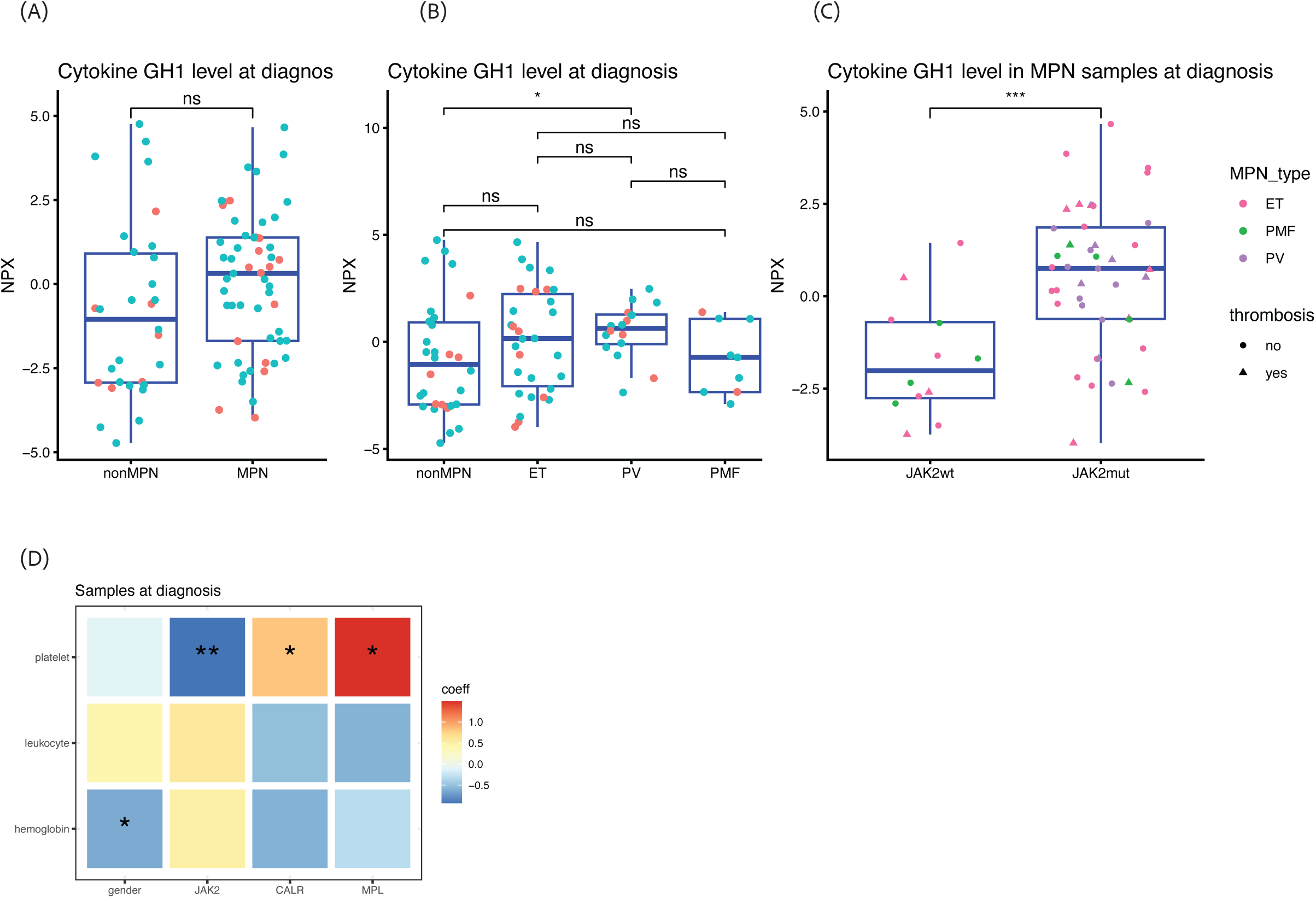
Boxplots of GH1 plasma level in MPN and non-MPN controls **(A)**, as well as in MPN subtypes (**B**); and in *JAK2*-mutant and wildtype MPN patients **(C).** (D) Associations between clinical variables, driver mutation status, and peripheral blood counts at diagnosis.

**Supplementary Figure 6.**
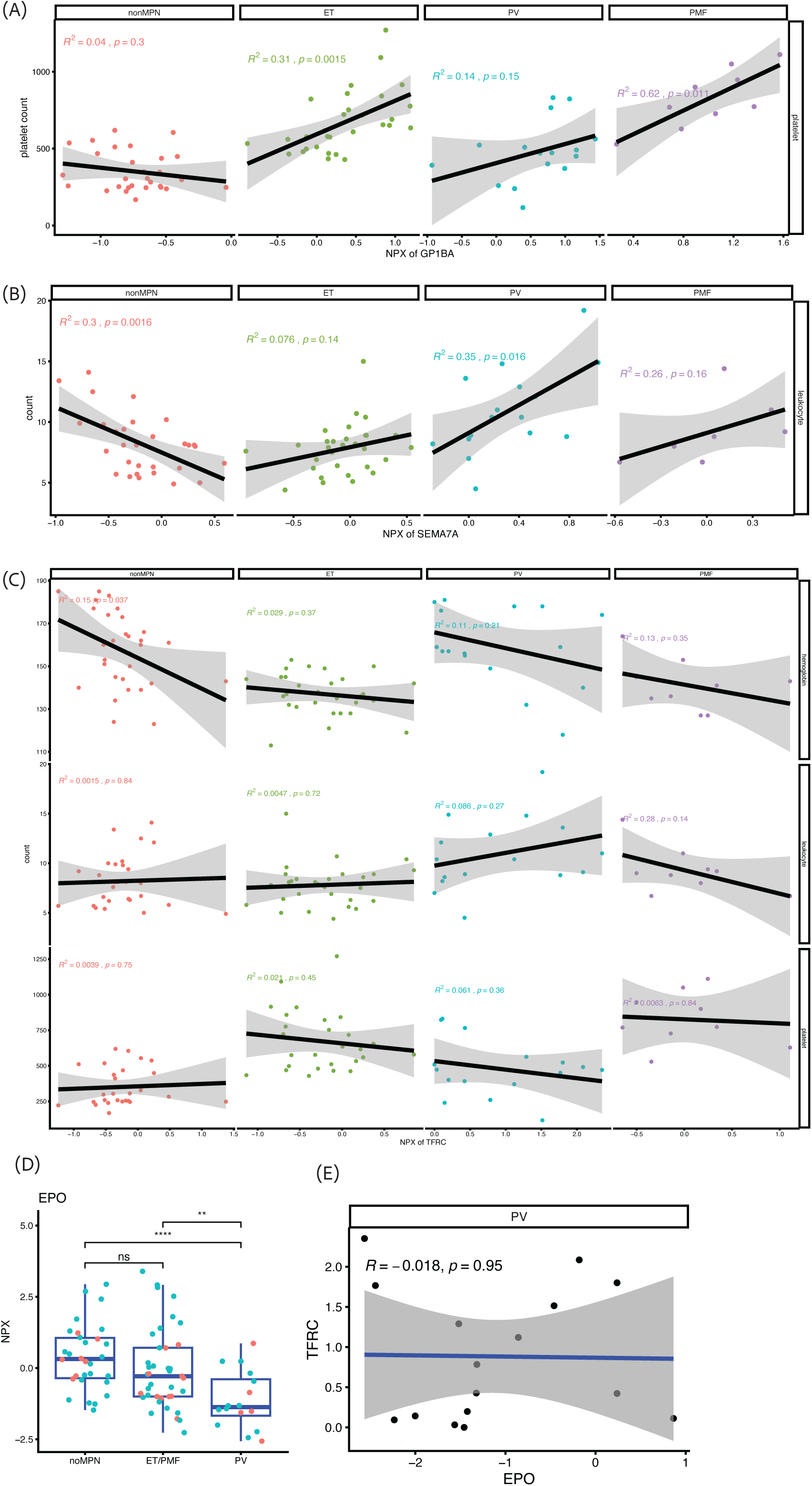
(**A**) Scatter plot showing correlation between GP1BA and platelet count across all patient groups at diagnosis. (**B**) Scatter plot showing correlation between SEMA7A and leukocyte count across all patient groups at diagnosis. (**C**) Scatter plot showing correlation between TFRC and hemoglobin/leukocyte/platelet count across all patient groups at diagnosis. (**D**) Boxplots showing EPO plasma protein expression level in non-MPN controls, ET/PMF and PV. Pairwise comparisons were performed using t-test. Significance levels: ns: p > 0.05; *: p <= 0.05; **: p <= 0.01; ***: p <= 0.001; ****: p <= 0.0001. (**E**) Linear regression between EPO and TFRC in PV samples at diagnosis.

